# MAX2-dependent signaling regulates the transition from 2D to 3D growth by suppressing cytokinin accumulation in *Physcomitrium patens*

**DOI:** 10.1101/2025.05.15.654175

**Authors:** Yi Luo, Yuki Hata, Juri Ohtsuka, Aino Komatsu, Mikiko Kojima, Hitoshi sakakibara, Junko Kyozuka

## Abstract

Plants undergo distinct developmental phases throughout their life cycle. Proper regulation of the timing of phase transition in response to internal and external cues is crucial to optimize proliferation and reproduction. In *Physcomitrium patens* (*P. patens*), a moss, the transition from growth of filamentous tissue growth called protonema to gametophore bud formation represents a shift from two-dimensional (2D) to three-demensional (3D) growrh, ultimately leading to reproductive development. MORE AXILLARY GROWTH2 (MAX2), an F-box protein, targets repressor proteins in the KARRIKIN INSENSITIVE2 (KAI2)-dependent signaling pathway for degradation via the 26S proteasome-dependent pathway. KAI2 perceives an unidentified ligand known as KAI2 ligand (KL) as well as karrikins. The four SMAX1-LIKE (SMXL) family proteins work as suppressors in the MAX2-dependent signaling in *P. patens*. Gametophore bud formation was accelerated when MAX2-dependent signaling was disrupted, while it was delayed in *Ppsmxl*s, loss of function mutants of the *PpSMXL*s. These results indicate that MAX2-dependent signaling suppresses gametophore bud formation in *P. patens*, which has not been previously described. We demonstrate that cytokinin activity and contents are increased in *Ppmax2* mutants, and cytokinin application rescued the delay in bud formation in *Ppsmxl* mutants, suggesting that MAX2-dependent signaling suppresses cytokinin accumulation. The four *SMXL* genes redundantly act to promote gametophore bud formation; PpSMXLC and PpSMXLD, which contain the conserved degron sequence, act as the repressor of MAX2-dependent signaling, while PpSMXLA and PpSMXLB function independent of proteolysis-dependent signaling. Based on these findings, we propose that modulating cytokinin activity to control developmental phase transition may be a conserved role of MAX2-dependent signaling in bryophytes.

**Significance Statement:** Plant hormones, both individually and in combination, regulate almost all aspects of plant growth and development. Gametophore bud formation in *Physcomitrium patens*, a moss, marks the transition from two-dimensional (2D) filamentous growth to three-dimensional (3D) shoot growth. We demonstrate that gametophore bud formation is controlled by the sequential action of two plant hormones, KL, an unidentified hormone that is transduced through MAX2-dependent signaling, and cytokinin. Four *SMXL* genes, repressors in MAX2-dependent signaling, redundantly promote bud formation: two function within the signaling pathway, while the other two operate independently of it. Our findings uncover novel mechanisms that directly link MAX2-dependent signaling to developmental phase transition by regulating stem cell activity, achieved through the regulation of cytokinin accumulation in *Physcomitrium patens*.

## Introduction

Plants undergo distinct developmental phases throughout their life cycle (1). Proper regulation of the timing of phase transition in response to internal and external cues is crucial to optimize proliferation and reproduction. The life cycle of *Physcomitrium patens* (*P. patens*), a moss, starts with the germination of a spore, followed by the planner growth producing filamentous tissues called protonemata, which are classified into chloronemata and caulonemata (2). The germinated spore produces chloronemal cells and develops chrolonemata. As development proceeds or the surrounding environment changes, caulonemal cells initiate. Caulonemata initiate gametophore branch initials, which differentiate into gametophore (leafy shoot) buds, in addition to the chrolonemal and caulonemal initials (3). Mature gametophores generate male and female reproductive organs and produce diploid sporophytes after fertilization. The diploid sporophytes produce new spores via meiosis and start the next generation. Among the distinct phase changes during the life cycle, changing from two-dimensional (2D) growth to three-dimensional (3D) growth is achieved by gametophore bud formation. The timing of this shift is crucial for the effective propagation of *P. patens* because this eventually leads to reproductive growth. Plant hormones play roles in the control of developmental phase changes in *P. patens*. Auxin regulates the transition from chloronema to caulonema formation (4). It is well known that cytokinins promote gametophore bud formation, however, the precise mechanisms by which cytokinins regulate bud formation remains unclear (5). Cell divisions of an apical cell of the gametophore that give rise to a bud meristem is trigged by cytokinins (6). Recently, we demonstrated that cytokinin levels increase specifically in the apical cell by upregulation of *LONEY GUY* (*LOG*) orthologs encoding the final enzyme for active cytokinin generation, during the gametophore formation (7–9). We also showed that cytokinin excludes PpTAWAWA, a putative differentiation factor, from the apical cell (8)

*KARIKIN INSENSITIVE 2* (*KAI2*), encoding an α/β hydrolase protein was identified as a receptor of karrikins, the chemicals contained in the burned plants (10). Subsequently, genetic analysis showed that KAI2 works as a receptor of an as-yet-identified endogenous molecule, which is tentatively called the KAI2 ligand (KL). *DWARF14* (*D14*), the strigolactone (SL) receptor, arose from the duplication of *KAI2* in the common ancestor of seed plants (11). In seed plants, the KL and SL signals are transmitted through partially overlapped proteolysis-dependent signaling pathways. Perception of KL and SL by KAI2 and D14, respectively, triggers the degradation of SUPPRESSORS OF MAX2 1 (SMAX1) and SMAX1-LIKE (SMXL) family proteins via MORE AXILLARY GROWTH2 (MAX2)-dependent ubiquitination (12–14). SMAX1/SMXL proteins are large multidomain protein structures that are generally highly conserved in land plants. In particular, RGKT, the degron motif in the D2 domain is well-conserved in the SMXL family proteins (15–17). In addition, Double Clp domain, which contains a nuclear localization signal, two NTPase domains, and the ethylene responsive element binding factor–associated amphiphilic repression (EAR) motif, which is required for interaction with transcriptional repressors, are well conserved (15–20). It was reported that SMXLs interact with TOPLESS to repress the transcription of downstream genes. Moreover, Recent research showed that SMXL6 directly binds DNA (21).

MAX2 is shared by KL and SL signaling pathways, while distinct sets of SMXL family proteins work as repressors in KL and SL signaling pathways. The *SMXL* genes form a small gene family (13, 17–19). In Arabidopsis, SMAX1 and SMXL2 are involved in the KAI2-dependent pathway that controls seed germination and hypocotyl elongation (12, 17). SMXL6, SMXL7, and SMXL8 are involved in the control of shoot branching regulated by SL signaling (13). SMXL3, SMXL4, and SMXL5 lost the sequence required for degradation during evolution. Therefore, they are not subjected to proteosome-dependent degradation (22).

Phylogenetic and genetic analysis showed that KL and SL signaling evolved step by step (18, 23–25). *D14*, the SL receptor, arose from the duplication of *KAI2* in the common ancestor of seed plants (11). *MAX2* gene is present from some algae species. On the other hand, SMXL proteins emerged from bryophytes. Therefore, the KAI2-dependent signaling was established during plant terrestrialization. The KAI2-dependent signaling regulates vegetative reproduction and dormancy of asexual buds via the degradation of MpSMXL in *Marchantia polymorpha* (*M. polymorpha*), a liverwort (26)*. P. patens* contains 13 *KAI2* family genes (*PpKAI2L*s) and one *MAX2* gene (27). *Ppmax2* mutant produces smaller protonema colonies and large gametophores (28, 29). This implies that *PpMAX2* promotes the protonemal growth. Recent studies performed using physiological and biochemical approaches revealed that some of the 13 PpKAI2Ls are involved in the MAX2-independent putative SL signaling. On the other hand, other PpKAI2Ls work in the KL signaling, which is MAX2-dependent (27, 28). *P. patens* contain four *SMXL* genes, *PpSMXLA* to *PpSMXLD* (29). It was reported that the four genes work in MAX2-dependent signaling. However, defects in complete loss of function mutants have not been described. Here, we show that the four genes redundantly act to promote gametophore bud formation by modulating cytokinin activity, a novel function.

## Results

### PpSMXLC and PpSMXLD, but not PpSMXLA and PpSMXLB, are subjected to MAX2-dependent protein degradation

The four *SMXL* orthologues of *P. patens*, Pp3c2_14220, Pp3c1_23530, Pp3c9_16100, and Pp3c15_16120 are designated as *PpSMXLA*, *PpSMXLB*, *PpSMXLC*, and *PpSMXLD*, respectively (Fig. 1*A*) (29). Phylogenetic analysis revealed that all four *PpSMXL*s genes belong to the same clade as the *SMXL* orthologues of *M. polymorpha* and are divided into two groups: *PpSMXLA* and *PpSMXLB* (designated as *PpSMXLA/B* hereafter) and *PpSMXLC* and *PpSMXLD* (designated as *PpSMXLC/D* hereafter), derived from a recent whole genome duplication (30). Among the four PpSMXLs, PpSMXLC and PpSMXLD show a high sequence similarity to MpSMXL and SMXL family proteins in angiosperms throughout the coding region. In contrast, PpSMXLA and PpSMXLB lack an NTPase I domain, whose function is unknown (Fig. 1*B* and *SI Appendix*, Fig. S1*A*). Moreover, the RGKT motif, which is required for the degradation of SMXL proteins (16), is changed to RGRT in PpSMXLA and PpSMXLB, making their proteolysis-dependent regulation questionable (*SI Appendix*, Fig. S1*A*). The EAR motif is conserved in all four PpSMXL proteins, suggesting that transcriptional suppression may be a common function of the four PpSMXLs.

**Fig. 1.**
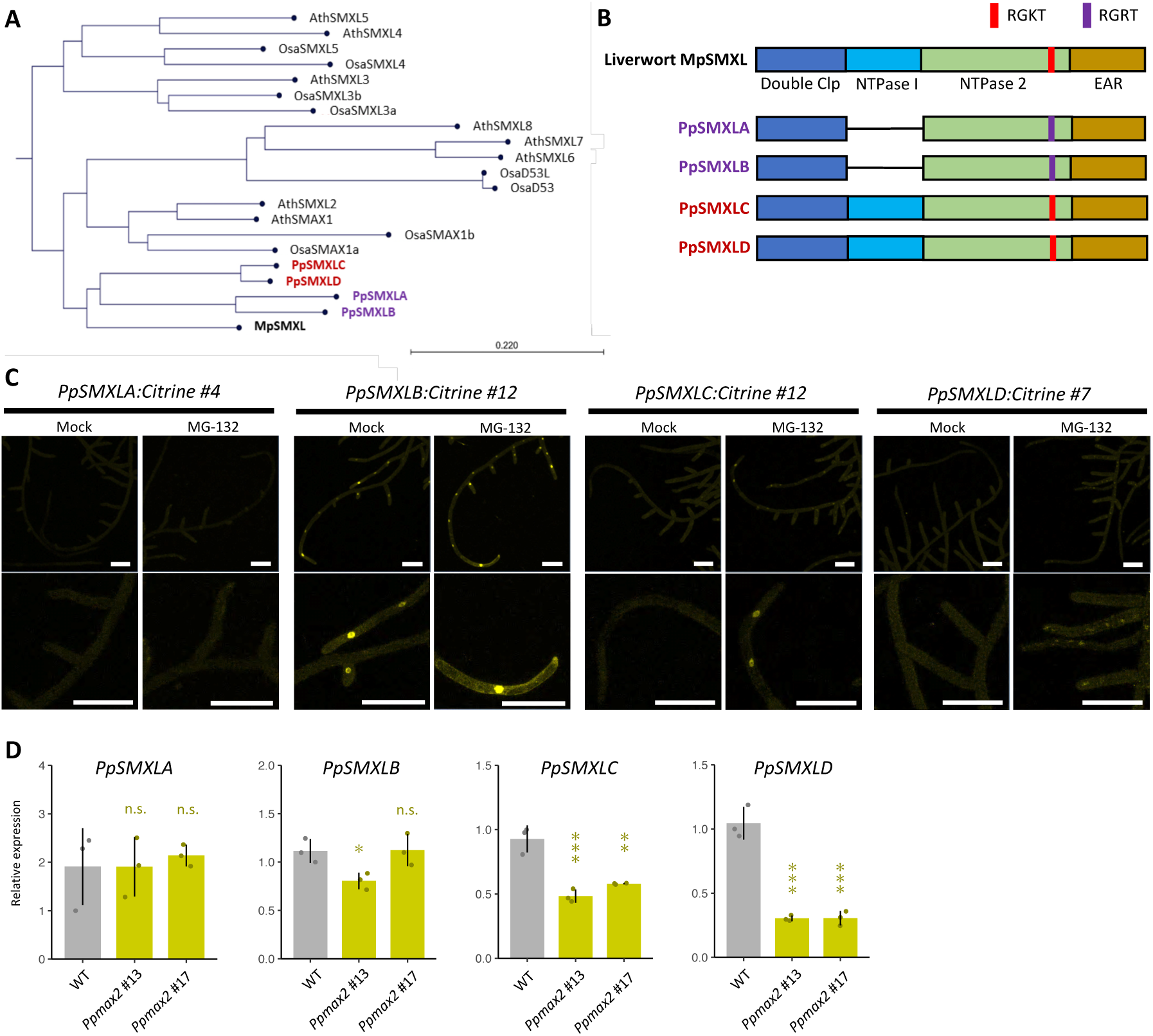
Characteristic analysis of four PpSMXLs. (**A**) Phylogenetic tree of SMXL family in land plants. Species abbreviations are as follows: *Marchantia polymorpha* (Mp), *Physcomitrium patens* (Pp), *Arabidopsis thaliana* (Ath), *Oryza sativa* (Osa). (**B**) Structures of PpSMXLs and MpSMXL proteins. (**C**) Localization of PpSMXL-Citrine fluorescence (yellow) in protonema treated with MG132 or control condition (Mock). Lower panels show tip region of a protonema. (**D**) Expression level of *PpSMXL*s in *Ppmax2* mutants and WT measured by qRT-PCR. Statistical significance was evaluated by Dunnett’s multiple comparison test between WT and mutants (n = 3, *p < 0.05, **p < 0.01, ***p < 0.001). Scale bars, 100 µm.

We tested the possibility that the four PpSMXLs are subjected to proteolysis-dependent regulation. A *Citrine* gene was knocked in the C terminus of the four *PpSMXL* genes by homologous recombination. In normal conditions, Citrine fluorescence was detected in nuclei of protonema cells in PpSMXLB:Citrine lines, but not in PpSMXLA:Citrine, PpSMXLC:Citrine, or PpSMXLD:Citrine lines (Fig. 1*C*). After the treatment of MG132, a proteasome inhibitor, Citrine fluorescence became visible in PpSMXLC:Citrine and PpSMXLD:Citrine lines. However, no signal was observed in PpSMXLA:Citrine lines even after MG132 treatment. In PpSMXLB:Citrine lines, apparent differences in the Citrine fluorescence intensity were not detected after the MG132 treatment. These results suggested that PpSMXLC and PpSMXLD, but not PpSMXLA and PpSMXLB, may be subjected to proteosome-dependent degradation (Fig.1*C* and *SI Appendix*, Fig. S1*B*). Despite the high expression of PpSMXLB protein, the transcriptional level of *PpSMXLB* is lower than that of *PpSMXLC* and *PpSMXLD*, suggesting that high accumulation of PpSMXLB:Citrine protein may be the consequence of escape from proteolysis (*SI Appendix*, Fig. S1*C*). Based on the increase in protein level in response to MG132 treatment, we assumed that PpSMXLC and PpSMXLD act as repressor proteins in the PpMAX2-dependent signaling pathway.

In the MAX2-dependent signaling pathway, the expression of *SMXL* genes is positively regulated by the signaling, a feedback mechanism thought to maintain the homeostasis of this pathway. This regulatory mechanism is well-conserved across a wide range of plant species, including angiosperms and *M. polymorpha* (31). To clarify whether the four *PpSMXL* genes are also influenced by PpMAX2-dependent signaling, we generated loss-of-function mutants of the *PpMAX2* gene and examined the expression of *PpSMXLA* to *PpSMXLD* (Fig. 1*D*). The expression levels of *PpSMXLA* and *PpSMXLB* were comparable to those in wild-type (WT). In contrast, the expression levels of *PpSMXLC* and *PpSMXLD* were decreased in *Ppmax2* mutants. These findings further support that PpSMXLC and PpSMXLD, but not PpSMXLA and PpSMXLB, are subjected to degradation in the MAX2-dependent signaling pathway and function as the repressor of the signaling pathway.

### PpMAX2-dependent signaling positively regulates colony size and cell size

To investigate the functions of *PpSMXL* genes, we generated degradation-resistant versions of PpSMXL proteins. We used *PpSMXLA* and *PpSMXLC* genes as the representatives of *PpSMXLA/B* and *PpSMXLC/D*, respectively. *PpSMXLA^d53^* and *PpSMXLC^d53^*, which lack the putative degron sequence (RGRT in PpSMXLA and RGKT in PpSMXLC), concatenated with β-estradiol inducible system were introduced into WT plants. We confirmed the β-estradiol-induced expression of *PpSMXLA^d53^* and *PpSMXLC^d53^* (*SI Appendix*, Fig. S2). We assumed that if *PpSMXLC* is involved in the PpMAX2-dependent proteolysis-based signaling, induction of *PpSMXLC^d53^-ox* would confer defects similar to that of *Ppmax2*. As previously reported, the colony size was significantly reduced in *Ppmax2* mutants (Fig. 2*A* and *B*) (28, 29). Although the cell length was reduced considerably in *Ppmax2,* the reduction does not seem to be sufficient to wholly account for the decrease in the colony size, suggesting that both cell elongation and cell division are likely affected in *Ppmax2* mutants (Fig. 2*C*). *PpSMXLC ^d53^-ox* showed defects resembling *Ppmax2*. In contrast, the colony size and cell length of *PpSMXLA^d53^-ox* were rarely affected compared to WT (Fig. 2*A* to *C*). These results support the notion that the MAX2-dependent signaling controls cell and colony extension through PpSMXLC and very unlikely PpSMXLA degradation.

**Fig. 2.**
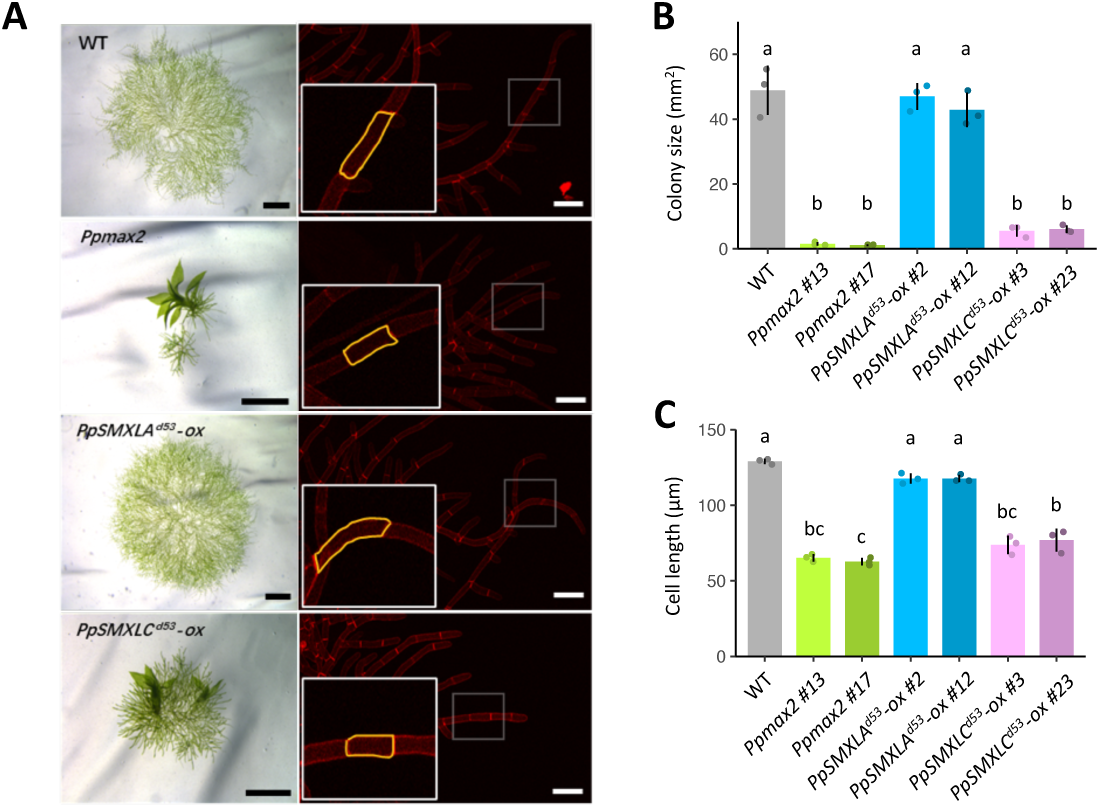
Plant colony size and protonema cell size in MAX2-dependent signaling mutants. (A) Plant colony of WT, *Ppmax2* mutants, and overexpression lines of *PpSMXLA^d53^* or *PpSMXLC^d53^*grown on BCD medium between two cellophane membranes for 17 days. In *PpSMXLA^d53^-ox* and *PpSMXLC^d53^-ox*, 1 nM β-estradiol was supplemented in the medium. Left panels show bright field images, and right panels show confocal microscopy images. In the confocal images, cell walls were stained with Propidium Iodide. Outlines of a cell is indicated by yellow lines. (**B** and **C**) Colony area (B) and length of protonema cells (C) in WT, *Ppmax2* mutants, and overexpression lines of *PpSMXLA^d53^* or *PpSMXLC^d53^*. The length of protonrmal cells was quantified by measuring the length of ten caulonema cells from the 2^nd^ cell on the edge to the 11^th^ cell and taking the average of them. Tukey’s honestly significant difference (HSD) test was used for multiple comparisons (n=3, p < 0.05, statistical differences are indicated by different letters). Black scale bars, 1 mm, and white scale bars, 100 μm.

### Gametophore bud formation is accelerated in *Ppmax2* and *PpSMXLC^d53^-ox* while delayed in *Ppsmxl* loss-of-function mutants

Previous studies reported increased gametophore size and altered protonema growth in *Ppmax2* mutants (28, 29). In this study, we recognized that the gametophore formation was accelerated in *Ppmax2*. Gametophores were first observed at 17 days after the inoculation (DAI) of protonemata cells in WT, while they were already formed 7 DAI in *Ppmax2*, indicating that the phase shift from protonema formation to gametophore formation was accelerated in *Ppmax2* (Fig. 3*A* and *B*). A similar extent of acceleration of gametophore bud formation occurred in *PpSMXLC ^d53^-ox* plants. The timing of gametophore bud formation was also accelerated in *PpSMXLA ^d53^-ox* compared to WT, while the effect was much weaker than that in *PpSMXLC ^d53^-ox* (15 DAI in *PpSMXLA ^d53^-ox* and 17 DAI in WT). These data indicate that the perturbation of PpMAX2-dependent degradation of PpSMXLC/D leads to earlier phase change.

**Fig. 3.**
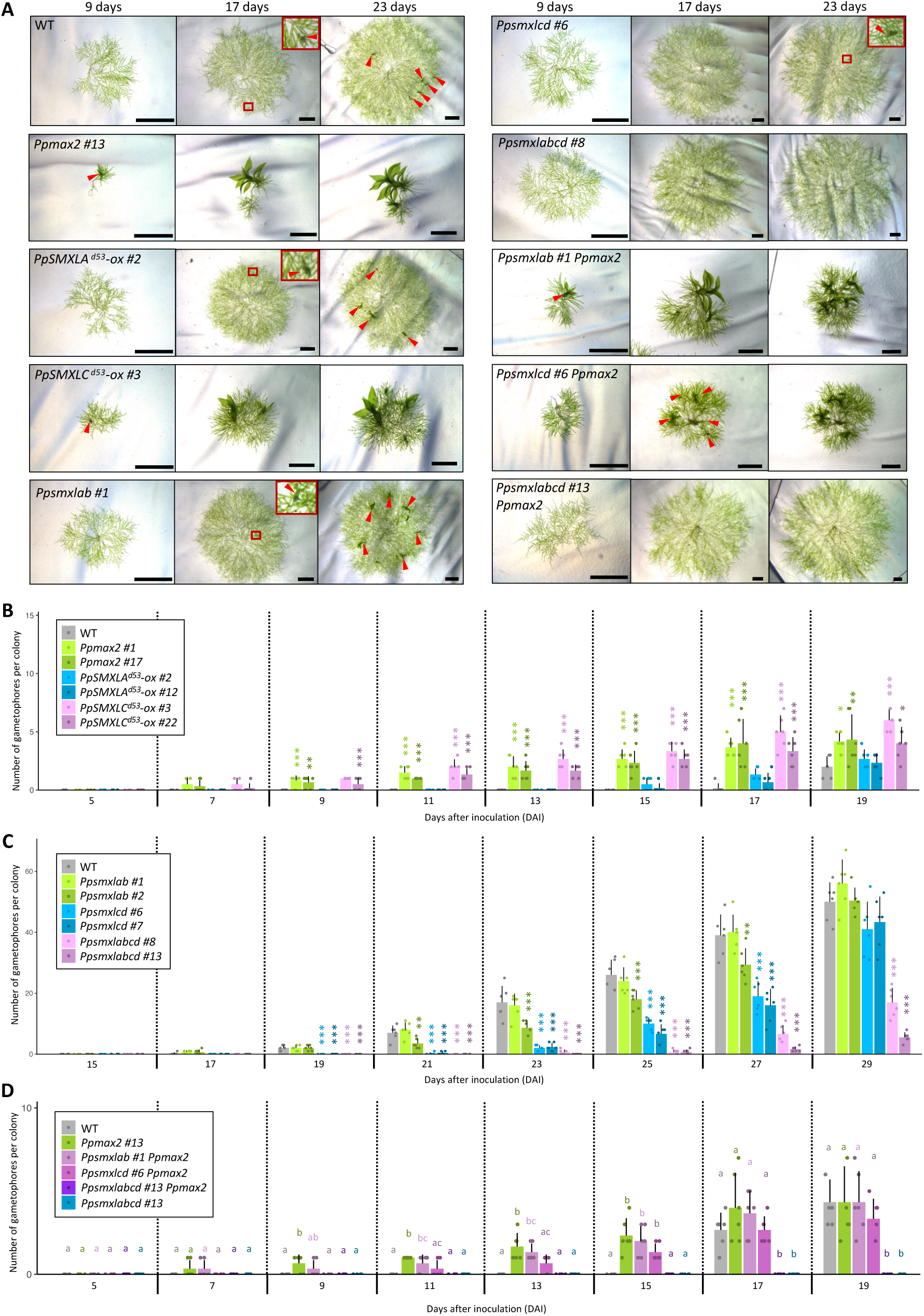
Phase change from protonema to bud formation in MAX2-dependent signaling mutants. (**A**) Plant colony of WT and MAX2-dependent signaling mutants grown on BCD medium between two cellophane membranes for 9, 17, or 23 days. In *PpSMXLA^d53^-ox* and *PpSMXLC^d53^-ox*, 1 nM β-estradiol was supplemented in the medium. Red arrowheads indicate gametophores. (**B** to **D**) Number of gametophores per colony in the WT and MAX2-dependent signaling mutants. Statistical significance was evaluated by Dunnett’s multiple comparison test between WT and mutants (n=6, *p < 0.05, **p < 0.01, ***p < 0.001), or HSD test (n=6, p < 0.05, statistical differences are indicated by different letters). Scale bars, 1mm.

We next generated *Ppsmxlab* (double mutants of *PpSMXLA* and *PpSMXLB*), *Ppsmxlcd* (double mutants of *PpSMXLC* and *PpSMXLD*), and *Ppsmxlabcd* (quadruple mutants of *PpSMXLA*, *PpSMXLB*, *PpSMXLC* and *PpSMXLD*) loss of function mutants by CRISPR/Cas9. The absence of the repressors in the signaling pathway is expected to allow maximum signaling. Gametophore bud formation was delayed to 21 DAI in *Ppsmxlcd* double loss-of-function mutants (Fig. 3*A* and *C*), indicating that PpSMXLC/D inhibit the function of suppressors of the phase transition in the MAX2-dependent signaling. Thus, the degradation of PpSMXLC/D or the loss of their functions leads to the suppression of gametophore bud formation, resulting in the delay of the phase change. Although the timing of the bud formation was not significantly affected in *Ppsmxlab*, it was much delayed in Ppsmxl*abcd* quadruple mutants (25 DAI). These findings suggest the involvement of *PpSMXLA/B* in controlling the phase change to gametophore formation. Since MG132 application experiments and overexpression of the degradation-resistant version showed no sign of the participation of PpSMXLA/B in the proteolysis-based signaling pathway, the function of PpMAX2A/B is very likely independent from PpMAX2 (Fig. 3*A* and *B*). Gametophores were eventually formed even in the *Ppsmxlabcd* quadruple mutants, implying that *PpSMXLA* to *PpSMXLD* genes control the timing of the phase change, not the process of bud formation itself.

We performed genetic analyses to confirm that PpSMXLC/D work in the same signaling pathway as PpMAX2. As described previously, buds were observed at 7 DAI in *Ppmax2*. The timing of bud formation in *Ppmax2* mutants was not significantly influenced by adding *Ppsmxlab* mutations, while combining *Ppsmxlcd* mutation partially rescued *Ppmax2*, supporting that PpSMXLC/D but not PpSMXLA/B work downstream of PpMAX2. Furthermore, combining the mutations of all four genes wholly rescued the precocious bud formation defects in *Ppmax2*, indicating the redundancy of PpSMXLA/B and PpSMXLC/D (Fig. 3*A* and *D*).

Based on the results of the proteolysis inhibitor application experiment, overexpression of degradation-resistant version repressors, defects in the loss of function mutants, and genetic analysis of *PpMAX2* and the four *PpSMXL* genes, we propose that *PpSMXLC/D* work in the PpMAX2-dependent signaling pathway and *PpSMXLA/B* work redundantly with *PpSMXLC/D*, but independently from PpMAX2 to suppress gametophore bud formation.

### PpMAX2-dependent signaling suppresses cytokinin accumulation

Gametophore bud formation in *P. patens* is promoted by cytokinin (5, 32). Therefore, we tested the possibility that the PpMAX2-dependent signaling regulates the timing of gametophore bud formation by controlling cytokinin function. To monitor cytokinin signaling, we used *TCSv2:GUS* lines in which the *GUS* gene is expressed under the control of TCSv2, a cytokinin-responsive regulatory sequence (8). We used two representative lines (*TCSv2:GUS* #11 and *TCSv2:GUS* #19) showing moderate expression levels and clear cytokinin responsiveness for further analysis. We mutated the *PpMAX2* gene of the selected two *TCSv2:GUS* lines by homologous recombination and compared the *TCSv2:GUS* expression levels with that in the WT background. We observed an evident upregulation of the GUS expression in the *Ppmax2* background in both TCSv2 marker lines, suggesting that PpMAX2 suppresses cytokinin activity (Fig. 4*A*). Next, we observed the spatial-temporal regulation of cytokinin signaling during colony growth. In WT, weak GUS signals were observed in the central part of the colony on DAI 5. The weak signal was expanded to the broader region except for the peripheral area of the colony by DAI 14. On DAI 21, the GUS signal became slightly more intense in caulonemata, and the buds showing the intense GUS signal appeared in WT (Fig. 4*B* and *SI Appendix*, Fig. S3*A*). In contrast, strong GUS signals were observed in the broad area of the *Ppmax2* colony on DAI 5. The signal became further intense when initiating buds that appeared on DAI 7. The pattern of GUS signal distribution was maintained throughout the culture time, with a slight decrease in the intensity in the central region at the later stage of colony development.

**Fig. 4.**
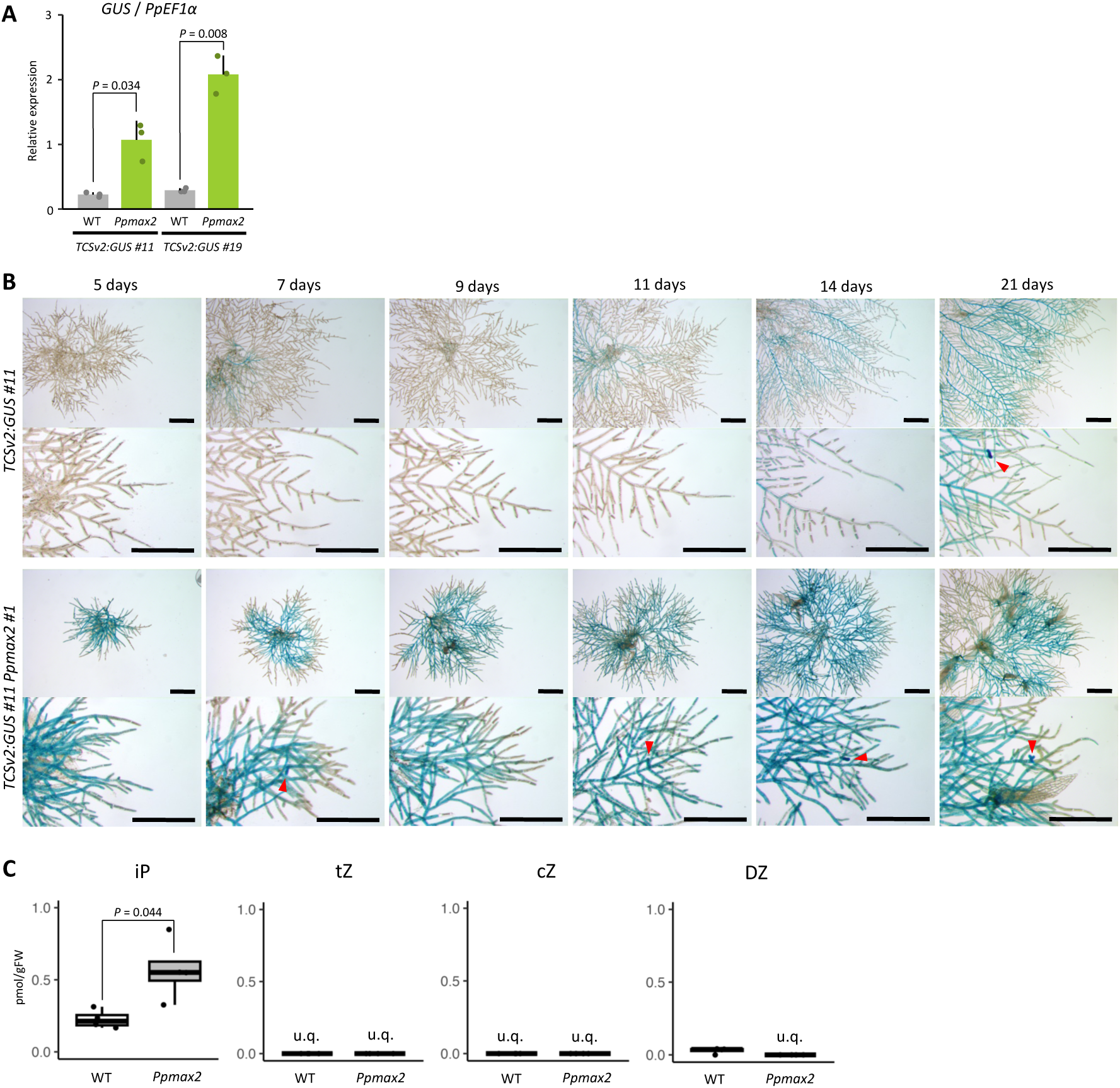
MAX2-dependent signaling regulates cytokinin level. (**A**) Expression level of *GUS* reporter genes driven by *TCSv2* cytokinin censor in WT and *Ppmax2* mutants. Statistical significance was evaluated by Student’s *t* test (n=3). (**B**) Expression patterns of *TSCv2:GUS* in WT and a *Ppmax2* mutant during plant colony growth. Lower panels in each genotype show tip region of protonemata. Red arrowheads indicate initiating buds. (**C**) Quantification of active cytokinin in WT and a *Ppmax2* mutant. u.q. indicates under quantification limits. Statistical significance was evaluated by Student’s *t* test (n=4). Scale bars, 500 µm.

So far, results indicate that PpMAX2-mediated signaling suppresses cytokinin signaling and/or cytokinin accumulation. To distinguish the two possibilities, we measured the cytokinin concentration. The level of *N^6^*-(Δ^2^-isopentenyl)-adenine (iP) was significantly increased in *Ppmax2* (Fig. 4*C* and *SI Appendix*, Fig. S3*B*). On the other hand, *trans*-zeatin (tZ), *cis*-zeatin (cZ), and dihydro-zeatin (DZ) were below the detectable level in both WT and *Ppmax2* except for low-level accumulation of DZ observed in WT. These results suggest that the PpMAX2-dependent signaling suppresses cytokinin accumulation.

### PpMAX2-dependent signaling delays gametophore bud formation through suppression of cytokinin accumulation

To support the hypothesis that the elevated cytokinin level in *Ppmax2* leads to the acceleration of bud formation, we introduced a gene encoding CYTOKININ OXIDASE (CKX), which catalyzes active cytokinins to inactive forms, to *Ppmax2* mutant (33). We assumed that *Ppmax2* defects would be alleviated in *Ppmax2*;*XVE:PpCKX1* lines in which cytokinin levels are lowered by the induction of *PpCKX1* expression. The early bud formation phenotype of *Ppmax2*;*XVE:PpCKX1* lines *w*as rescued by introducing *PpCXX1* expression by the addition of estrogen (Est+). Without induction of *PpCKX1* expression (Est-), many buds appeared by DAI 9. On the other hand, no bud was observed even at DAI 23 if *PpCKX1* was induced (Est+) (Fig. 5*A* and *B*). These results support the hypothesis that over-accumulation of cytokinin caused precocious bud formation in *Ppmax2*.

**Fig. 5.**
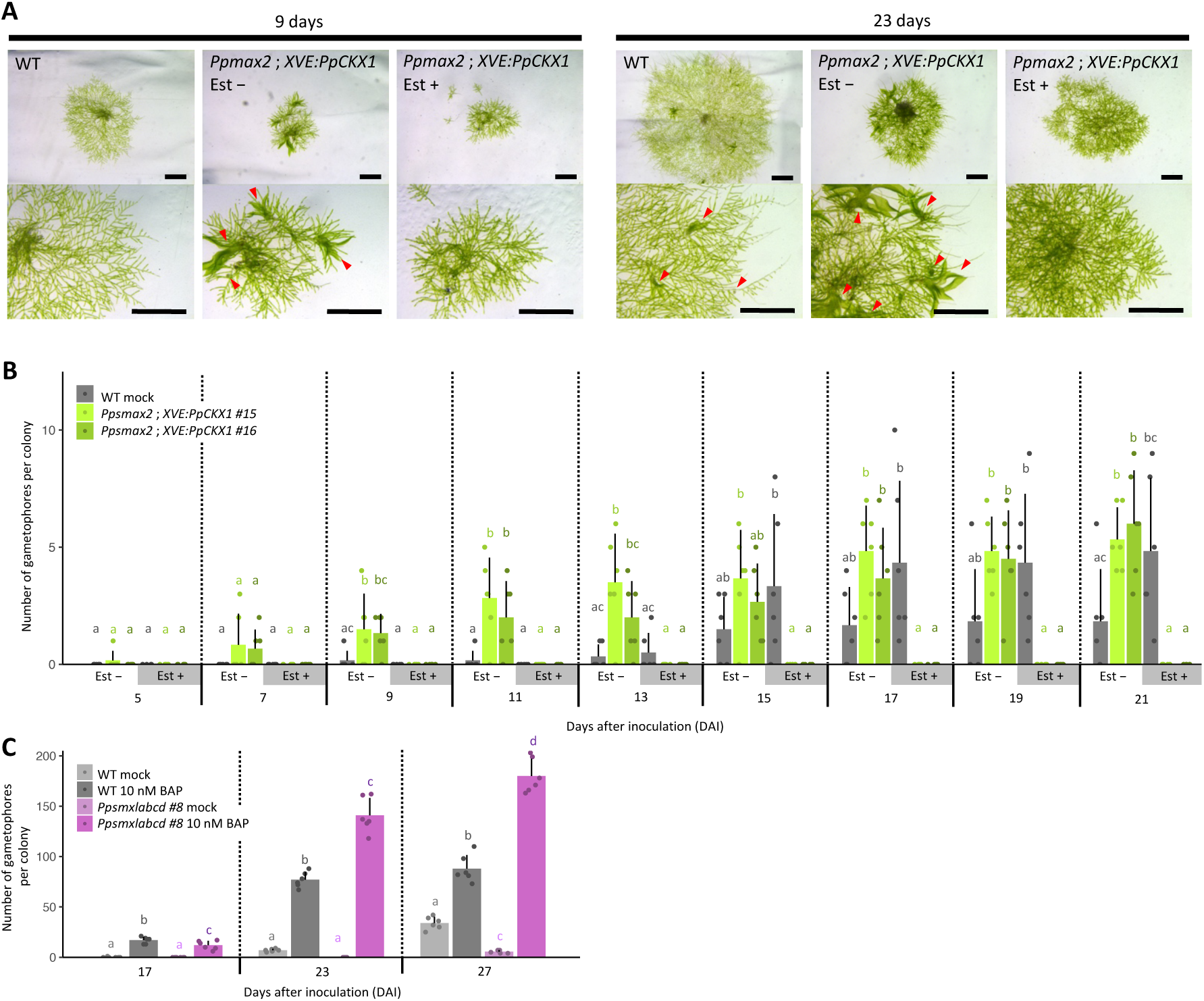
The gametophore formation regulated by MAX2-dependent signaling depends on cytokinin levels. (**A**) Plant colony of WT and *Ppsmax2;XVE:PpCKX1* lines grown between two cellophane membranes on BCD medium supplemented with 1 µM β-estradiol (Est +) or solvent (Est -). Lower panels show edge region of the colony. Red allowheads indicate gametophores. (**B**) Number of gametophores per colony in WT and *Ppsmax2;XVE:PpCKX1* grown on BCD medium supplemented with 1 µM β-estradiol (Est +) or solvent (Est -). Statistical significance was evaluated by HSD test (n=6, p < 0.05, statistical differences are indicated by different letters). (**C**) Number of gametophores per colony in WT and *Ppsmxlabcd* quadruple mutants grown on BCD medium supplemented with 10 nM BAP or solvent (Mock). Statistical significance was evaluated by HSD test (n=6, p < 0.05, statistical differences are indicated by different letters). Scale bars, 1 mm.

To further pursue this possibility, we tested if adding cytokinin rescues the defects in *Ppsmxlabcd* mutants. We analyzed the effects of the exogenous cytokinin supply on the number and timing of gametophore bud formation (Fig. 5*C* and *SI Appendix*, Fig. S4). In WT, buds were hardly observed on DAI 17 without BAP, while application of BAP accelerated bud formation in WT and *Ppsmxlabcd*. The number of buds per colony was slightly lower in *Ppsmxlabcd* than in WT at DAI 17, while more buds were formed in *Ppsmxlabcd* by DAI 23 than in WT when BAP was applied. Therefore, adding cytokinin wholly rescued bud formation defects in *Ppsmxlabcd* quadruple mutants, implying that PpSMXLA to D controls the timing of the growth phase change from protonemal growth to gametophore bud formation through positively controlling cytokinin accumulation (Fig. 5*C* and *SI Appendix*, Fig. S4). When the MAX2-dependent signaling is functional, degradation of PpSMXLC/D leads to lower cytokinin accumulation, resulting in the delay in bud formation. The number of buds was higher in *Ppsmxlabcd* than in WT when treated with BAP. We consider this probably to reflect the increased colony size in the quadruple mutants.

## Discussion

We demonstrated that MAX2-dependent signaling suppresses gametophore bud formation in *P. patens*, a function that has not been previously described. Bud formation was accelerated in *Ppmax2* mutants, while it was delayed in the loss of function mutants of *PpSMXL*s, which are suppressors of MAX2-dependent signaling. Cytokinin activity was increased in *Ppmax2*, *Ppmax2* defects were rescued by expressing CKX, and the application of cytokinin rescued the delayed bud formation in *Ppsmxl* mutants, suggesting that MAX2-dependent signaling suppresses bud formation by inhibiting cytokinin accumulation.

### MAX2-dependent signaling regulates the timing of growth phase change through cytokinin

When the MAX2-dependent signaling is acitve, degradation of PpSMXLC/D leads to the activation of factors repressed by PpSMXLC/D, resulting in the inhibition of cytokinin accumulation. Thus, factors repressed by PpSMXLC/D likely play a role in suppressing cytokinin accumulation. Given that PpSMXLC/D work as repressors of transcriptional factors (TFs) (19), the TFs likely control genes involved in inhibiting cytokinin accumulation (Fig. 6).

**Fig. 6.**
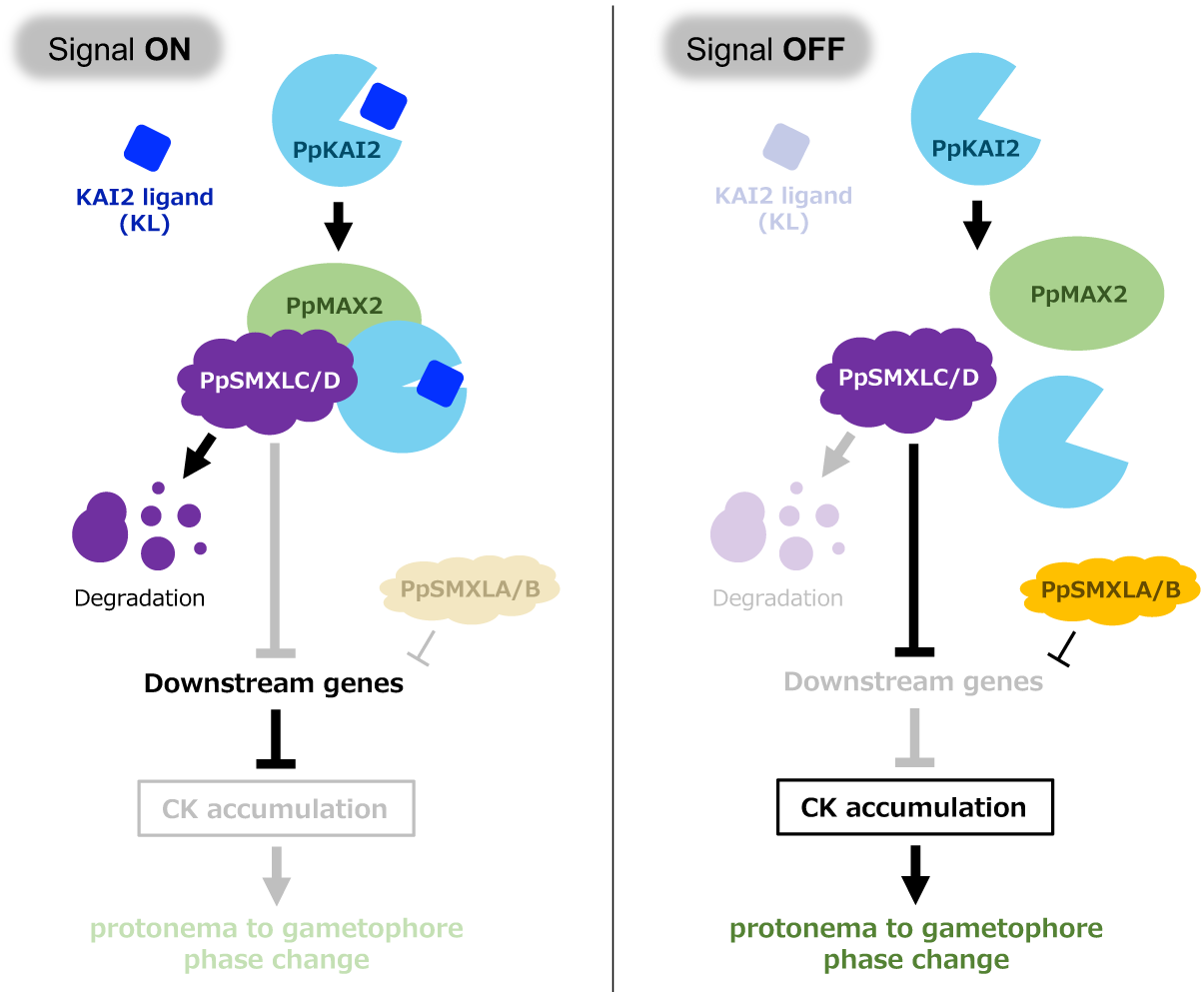
Protonema to gametophore phase change regulated by MAX2-dependent signaling in *P. patens*. When the signal pathway is active (Signal ON), the KAI2 ligand (KL) is perceived by the PpKAI2 and induces the formation of the complex with PpMAX2 and PpSMXLC/D, leading to the degradation of PpSMXLC/D, transcriptional repressors. Downstream genes inhibiting cytokinin (CK) accumulation are expressed. Thus, the gametophore formation does not occur, and the protonema phase continues. When the signal pathway is inactive (Signal OFF), the PpSMXLC/D are stable and repress the downstream genes inhibiting CK accumulation. It increases the CK level, leading to the protonema to gametophore phase change. PpSMXLA/B have redundant functions with the PpSMXLC/D, but are not regulated by the MAX2-dependent signaling. Expression levels of PpSMXLA/B are likely regulated by other mechanisms, such as transcriptional regulation.

Cytokinin is well-known to activate gametophore bud formation in *P. patens*. However, the regulation of cytokinin levels remains poorly understood. Recently, we demonstrated that cytokinin levels increase specifically in the apical initial cell during the gametophore bud formation (7, 8). Furthermore, a subset of the nine *LOG* orthologs in *P. patens* is specifically up-regulated in the gametophore apical cell. This suggests that LOG expression may contribute to the specific accumulation of cytokinin in the apical cell, thereby specifying its identity as the apical cell identity. This discovery positions *PpLOG*s as potential downstream components of the MAX2-dependent signaling pathway that regulates bud formation. In *M. polymorpha*, MAX2-dependent signaling controls vegetative reproduction by regulating cytokinin through the expression of Mp*LOG*, the single *LOG* gene in *M. polymorpha* (34). *LOG* homologs are found across major prokaryotic lineages of as well as plants, indicating an ancient origin (35). The control of cytokinin activity by regulating *LOG* expression may represent an ancestral function of MAX2-dependent signaling.

In addition to LOG, which catalyzes the final step of cytokinin biosynthesis, other components of the cytokinin biosynthesis pathway are also potential candidates for the factors controlled by MAX2-dependent signaling. However, less is known in bryophytes compared to the advances made in elucidating cytokinin synthesis pathways in angiosperms (36). In angiosperms, cytokinins are synthesized through adenosine phosphate-ISOPENTENYLTRANSFERASEs (IPTs) or tRNA-IPTs. In the former pathway, oxygenation of iP riboside 5’-phosphates (iPRPs) by CYP735A produces tZ riboside 5’-phosphates (tZRPs). LOG subsequently synthesizes active cytokinins from nucleotide-precursors, iP riboside 5’-monophosphate (iPRMP), tZ riboside 5’-monophosphate (tZRMP), DZ riboside 5’-monophosphate (DZRMP) and cZ riboside 5’-monophosphate (cZRMP) (37). Phylogenetic analysis showed that cognate orthologs of angiosperm adenosine phosphate-IPTs and CYP735A are absent in bryophytes. In contrast, *LOG* genes are well conserved in bryophytes. Further analysis of the cytokinin biosynthesis pathway is also required to elucidate the link between MAX2-dependent signaling and the control of bud formation. Besides biosynthesis, degradation is also a crucial step in adjusting hormone levels. Cytokinins are irreversibly deactivated by CKX (33). Overexpression of *PpCKX1* inhibits gametophore growth (38). *PpCKX1* is expressed in the apical cell and the surrounding cells in the gametophores, likely reflecting a feedback mechanism of cytokinin homeostasis in which C*KX* expression is activated by cytokinin (32). Therefore, we cannot rule out the possibility that MAX2-dependent signaling positively regulates *PpCKX*s and/or other regulators of cytokinin deactivation.

### Role of MAX2-dependent signaling in bryophytes

We demonstrated that MAX2-dependent signaling suppresses cytokinin accumulation in *P. patens*, while it enhances cytokinin contents in *M. polymorpha*. Thus, although the regulation of cytokinin accumulation by MAX2-dependent signaling is a common feature, its effects are opposite in the two bryophytes. In *P. patens*, suppression of cytokinin levels leads to delayed gametophore bud formation, whereas enhancement of cytokinin levels promotes vegetative reproduction in *M. polymorpha* (26, 31). Considering that gametophore formation ultimately leads to the first step of reproduction in *P. patens*, we can interpret the function of the MAX2-dependent signaling as maintaining the vegetative phase by modulating cytokinin levels in both *P. patens* and *M. polymorpha*.

MAX2-dependent signaling in bryophytes is thought to be triggered by perception of an unidentified endogenous ligand, referred to as KL. In *P. patens*, gametophore bud formation is influenced by light quality - enhanced by red light and inhibited by blue light (39) - suggesting that the KL signal may be regulated by light. Similarly, gemma and gemma cup formation in *Marchatia* species are also controlled by light quality (40, 41). In Arabidopsis, KL signaling controls seed germination, a trait controlled by light (14, 42). Therefore, the regulation of MAX2-dependent signaling by light may represent an ancestral function of this pathway, conserved across land plants. The next challenge is to understanding the environmental signals that regulate the activation and deactivation of MAX2-dependent signaling.

### Functional diversification of *SMXL* genes by gene duplication in *P. patens*

In general, components of plant hormone signalings pathways have increased in copy number through gene duplication during evolution (43). This is also the case for *SMXL* family genes, and the expansion of the *SMXL* gene family has extensively contributed to the diversification of the functions of KL and SL signaling pathways (18, 19). Among the eight SMXL proteins in Arabidopsis, five contain the RGKT motif, which is required for proteolysis, and are subjected to MAX2-dependent degradation for signaling of SL and KL (13, 17, 20, 21). In contrast, proteins from the SMXL4 subclade of Arabidopsis, namely SMXL3, SMXL4, and SMXL5, lack the RGKT motif and control root vasculature development independently of KL or SL signaling (19, 22). This suggests that the loss of the degron motif is an alternative mechanism for generating novel regulation, unaffected by hormonal signaling.

The results presented in this study suggest that, among the four *SMXL* genes in *P. patens*, *PpSMXLA/B* have a modified RGKT sequence, changed to RGRT, in addition to the loss of an NTPase domain of unknown function. These genes are independent of the proteolysis-dependent signaling. In contrast, *PpSMXLC/D*, which maintain a conserved domain structure similar to that of seed plans SMXLs, including the RGKT motif, function as repressors of MAX2-dependent signaling (20). Although PpSMXLA/B are unlikely to be subjected to the degradation, they play a redundant role with PpSMXLC/D in promoting bud formation. Phylogenetic analysis indicates that following the duplication of the ancestral genes for *PpSMXLA/B* and *PpSMXLC/D*, each pair underwent further duplication and divergence to form *PpSMXLA* and *PpSMXLB*, and *PpSMXLC* and *PpSMXLD*. The modification of the degron sequence occurred prior to the duplication of *PpSMXLA* and *PpSMXLB*. The very low levels of PpSMXLA expression suggest that, If they contribute to any function, it is to a lesser extent. In contrast, the PpSMXLB protein accumulates at higher levels. The absence of proteolysis may allow the accumulation of PpSMXLB protein, irrespective of signaling function. The diversification of the four *PpSMXL* genes may be an evolutional innovation to optimize growth, adapting to environmental conditions.

## Materials and Methods

### Plant materials and growth conditions

A wild-type strain derived from Gransden Wood (44) was used for transformation, phenotypic analysis, and gene expression analysis.

Plants were grown and propagated on BCDAT solid medium plates covered on cellophane at 25 °C under continuous light (45). For localization analysis of PpSMXLs-Citrine, a small amount of protonema filaments was taken and subcultured on BCD medium plates sandwiched between two cellophane membranes for 2 weeks. Then, the cultured protonema tissues were treated by 100 μM MG132 (26S proteasome inhibitor) or solvent (water containing 1% [v/v] DMSO) for 6 hours (26). For morphological observation and cytokinin quantification, colonies of WT and mutants were grown on BCDAT medium plates for 2 weeks, and then a small amount of protonema filaments on the edge of the colony was taken and subcultured on BCD medium sandwiched between cellophane membranes for 1-4 weeks. In drug treatment experiments, protonemata were grown on BCD medium containing β-estradiol (1 nM or 1 µM), 10 nM 6-benzylaminopurine (BAP, a synthetic CK) or solvent (water containing DMSO). For gene expression analysis by qRT-PCR, protonemata were grown on BCD medium plates covered on cellophane for 6 days. Then, in *PpSMXLA^d53^-ox*, *PpSMXLC^d53^-ox* or *XVE:PpCKX1* lines, protonemata were treated by 1 nM β-estradiol or solvent (water containing DMSO) for 6 hours.

### Construction of the alignment and Phylogenetic Tree

Orthologs of SMXL family sequences were collected by BlastP search using MpSMXL as a query on Phytozome (https://phytozome.jgi.doe.gov/pz/portal.html) and National Center of Biotechnology information (http://www.ncbi.nlm.nih.gov). MUSCLE alignment was performed by Geneious (ver. 10.2.6) and conserved P-Loop NTPase-domain and Double Clp-N motif domain sequences were used for phylogenetic analysis. The phylogenetic tree was constructed using the neighbor-joining method and bootstrap analysis (1,000 replicates) on the CLC sequence viewer (ver. 8.0).

### Vector contruction

The pCit-aphIV vector (46) was used for localization analysis of PpSMXLs. A DNA fragment containing genomic region coding C-terminus of PpSMXLs (stop codon was replaced with a linker sequence, GGAGGAGGATCA), and the genomic region of the 3’UTR of each *PpSMXL* gene (approximately 1.1-1.5 kb each), were amplified by PCR. These were cloned upstream or downstream of the Citrine coding region and antibiotic resistance gene cassette of the pCit-aphIV vector, respectively (*SI Appendix*, Fig. S5).

pTN186 vector (46) was used for gene disruption of *PpMAX2*. Approximately 1.2 kb each of the 5’UTR and 3’UTR of *PpMAX2* gene were amplified by PCR and cloned into the 5’ or 3’ sides of the antibiotic resistance gene cassette of pTN186, respectively (*SI Appendix*, Fig. S6).

pPGX8 vector (47) was used for β-estradiol-dependent overexpression of *PpSMXLA^d53^*, *PpSMXLC^d53^*, or *PpCKX1* (*SI Appendix*, Fig. S7). The protein coding regions of each gene were amplified by PCR using cDNA as a template and cloned into pENTR/D-TOPO vector (Invitrogen). The sequence encoding degron of PpSMXLA (AGAGGGCGTACA) or PpSMXLC (CGAGGTAAAACA) was deleted by inverse PCR in the pENTR/D-TOPO vector. The produced DNA fragment encoding PpSMXLA^d53^ or PpSMXLC^d53^ in pENTR/D-TOPO was transferred to pPGX8 vector by LR reaction. In case of *PpCKX1*, DNA fragment containing LexA operator and coding sequence of XVE was amplified by PCR and introduced to the EcoRV site of pTN182 to produce pPGX8-nptII. The DNA fragment encoding PpCKX1 in pENTR/D-TOPO was transferred to pPGX8-nptII by LR reaction.

Creating vectors for the guide RNA of the CRISPR/Cas9 system followed the protocol of Lopez-Obando et al. (48) (*SI Appendix*, Fig. S8*A*). The webtool CRISPOR (http://crispor.tefor.net/) was used for target sequence determination (*SI Appendix*, Fig. S8*B*). For the generation of double mutants (*Ppsmxlab*, *Ppsmxlcd*), and quadruple mutants (*Ppsmxlabcd*), we selected a 20 bp sequence a target of four *PpSMXL*s, which are near the start codon in the protein coding regions. To produce PpSMXLA, PpSMXLB, PpSMXLC and PpSMXLD sgRNA plasmids, a 500 bp fragment containing the snRNA U6 promoter followed by the sgRNA scaffold and flanked by attB recombinant sites, was chemically synthesized (Integrated DNA Technologies). The fragment was cloned into a pDONR207 backbone by the BP-reaction (49). Then, the target sequence for each PpSMXLs was introduced to the sgRNA by site-directed mutagenesis with inverse PCR. The introduced target sequence for each PpSMXLs was confirmed by Sanger sequencing.

All primers used for vector construction are listed in *SI Appendix*, Table S1.

### Generation of transgenic lines and mutants

Transformation of *P. patens* followed a protocol in (45). In short, protoplasts were made from protonemata cultured on BCDAT medium plates covered on cellophane membrane for 6 to 7 days, and the polyethylene glycol-mediated transformation was conducted. One μg pBNRf, 1 μg pAct-Cas9, and 5 μg pENTPp-U6-sgRNA were used for gene disruption by CRISPR/Cas9. For the introduction of pCit-aphIV and pPGX8 vectors, only the region to be integrated into the genome by homologous recombination was excised from each vector with appropriate restriction enzymes, and 15 to 25 µg of DNA fragments purified by ethanol precipitation were used. For the introduction of *PpMAX2* disruption construct by homologous recombination, only the region to be integrated into the genome was amplified by PCR, and 15 μg of each DNA fragment purified by ethanol precipitation was used.

For genotyping transformants by homologous recombination, PCR was performed using primers designed for the genomic regions inside and outside the introduced construct to confirm targeting (*SI Appendix*, Fig. S5*B*, S6*B*, and S7*B*). For genotyping transformants using CRISPR, genomic regions around the target sequence were amplified by PCR and mutations were confirmed by Sanger sequencing (*SI Appendix*, Fig. S8*C*).

All primers used for genotyping are listed in *SI Appendix*, Table S1.

### Histochemical GUS activity assay

GUS staining was performed following (46) with minor modifications. Plant tissues were fixed with a fixation solution [0.2% (w/v) MES (pH 5.6), 0.3% (v/v) formalin, 0.3 M mannitol] for 10 minutes at room temperature. After washing with 50 mM NaH_2_PO_4_ (pH 7.0), the fixed tissues were vacuum-infiltrated with a substrate solution [50 mM NaH_2_PO_4_ (pH 7.0), 0.5 mM 5-bromo-4-chloro-3-indolyl β-D-glucuronide (X-Gluc), 0.5 mM K_3_Fe(CN)_6_, 0.5 mM K_4_Fe(CN)_6_, and 0.05% (v/v) Triton X-100] for 30 minutes and then stained at 37°C. After staining, the tissues were fixed with 5% (v/v) formalin for 10 minutes and subsequently immersed in 5% (v/v) acetic acid for 10 minutes. The stained and fixed tissues were dehydrated with an ethanol series.

### Microscopy

Phenotypic analyses were performed using microscopic observation and digital images were captured using a Leica M165FC or Olympus BX51 with a UPlanFl20x/0.50 or UPlanFl40x/0.75 objective lens. For fluorescence observation, a confocal laser scanning microscope LSM880 (Carl Zeiss) was used. An LD LCI Plan-Apochromat 40x/1.2 Imm Korr DIC M27 or Plan-Apochromat 20x/0.80 M27 objective lens was used. For cell size measurement, the cell walls were stained with 1% Propidium Iodide (PI) before observation. Citrine was excited at 514 nm, whereas cell walls stained with PI were excited at 543 nm.

### RNA extraction

Plant tissues were collected from the medium, blotted with paper towels to absorb excess moisture, and then quickly frozen in liquid nitrogen. Frozen tissues were crushed with a multi-bead shocker (Yasui Kikai) and NucleoSpin RNA Plant (Macherey-Nagel) was used to extract RNA follwing the manufacture’s instructions.

### RT-qPCR analysis

cDNA was synthesized from total RNA using the SuperScript VILO cDNA Synthesis Kit (Invitrogen). Quantitative realtime PCR was performed using KOD SYBR qPCR Mix (TOYOBO) with LightCycler480II (Roche) or CFX Duet (BIO-RAD). *PpEF1α* (Pp3c2_10310) was used as a reference gene (49). Primers used in qRT-PCR are listed in Table S1.

### Quantification of cytokinins

Ninety colonies (∼100 mg fresh weight) of WT and *Ppmax2 #13* were freeze-dried using a freeze dryer (FDM-1000, EYELA, Japan). The freeze-dried samples were crushed, soaked in extraction solvent, and fractionated (50), and the semi-purified cytokinin fractions were subjected to mass spectrometric analysis using ultra-performance liquid chromatography (UPLC)-tandem mass spectrometry (AQUITY Premire UPLC System/XEVO-TQXS; Waters, USA) with an ODS column (ACQUITY Premier HSS T3 with VanGuard FIT, 1.8 µm, 2.1×100 mm; Waters, USA). All parameters, settings, and analytical procedures were determined following the methods described previously (34). Four biologically independent samples were measured and the mean values were calculated.

## Acknowledgments

This research was supported by a Grants-in-Aid from the Ministry of Education, Culture, Sports, Science, and Technology, Japan (23H05409, 20H05684, and 17H06475 to J.K., 20J20812 and 23K19362 to Y.H.).

