## Supplementary material for "MAX2-dependent signaling regulates the transition from 2D to 3D growth by suppressing cytokinin accumulation in *Physcomitrium patens*": SI Appendix

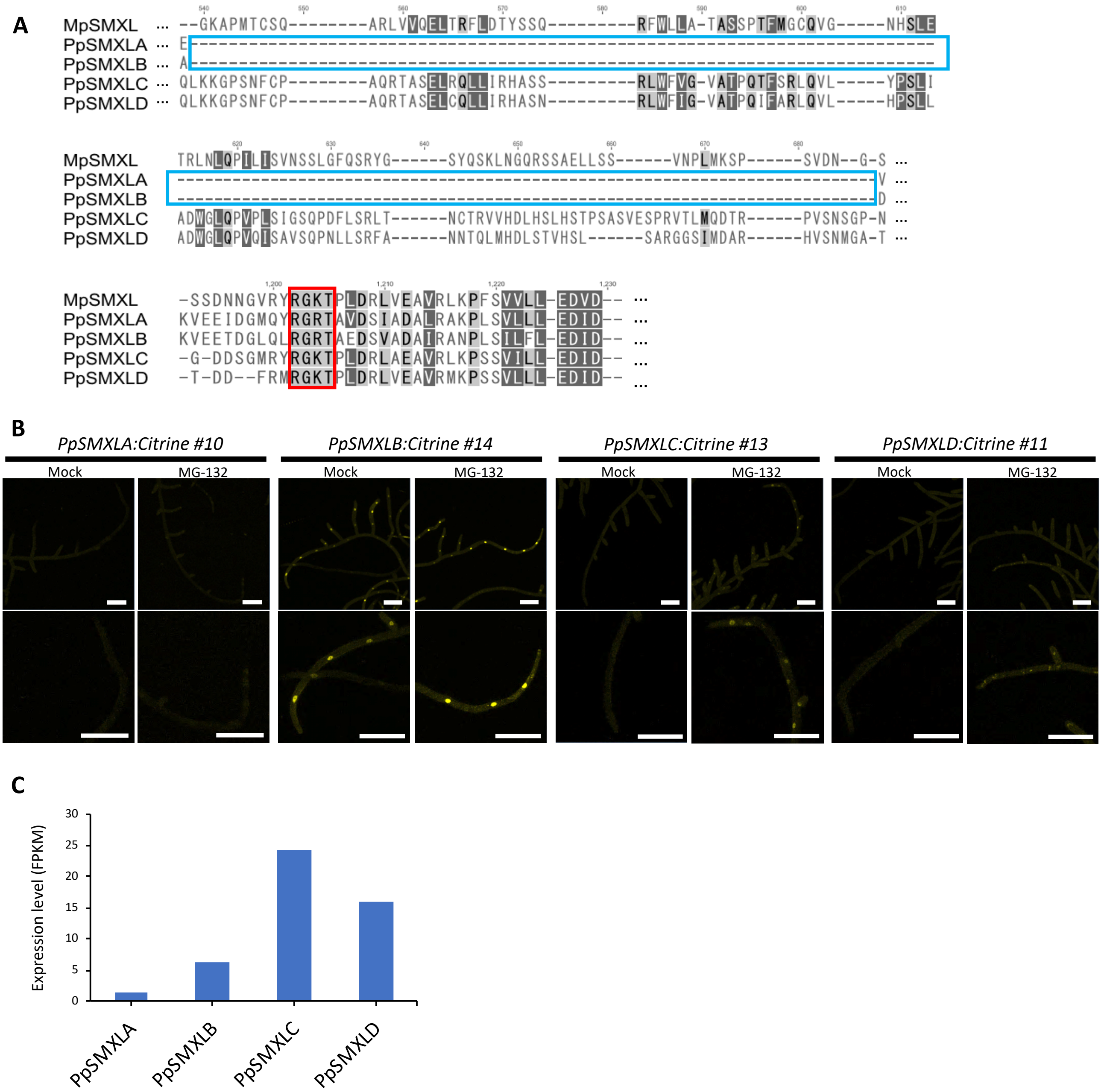


**Fig. S1. Domain structure and cellular localization of PpSMXL proteins.**

(A) Alignment of protein sequences in PpSMXLs and MpSMXL. Blue frames represent lack of NTPase I domain in PpSMXLA and PpSMXLB; Red frame represents RGKT motif. (B) Localization of PpSMXL-Citrine fluorescence (yellow) in protonema treated with MG132 or control condition (Mock). Lower panels show tip region of a protonema. (C) The expression level of *PpSMXL* genes in the WT protonema grown on the BCD solid medium. Data were obtained from public transcriptome data in the PEATmoss database (1). Scale bars, 100 μm.

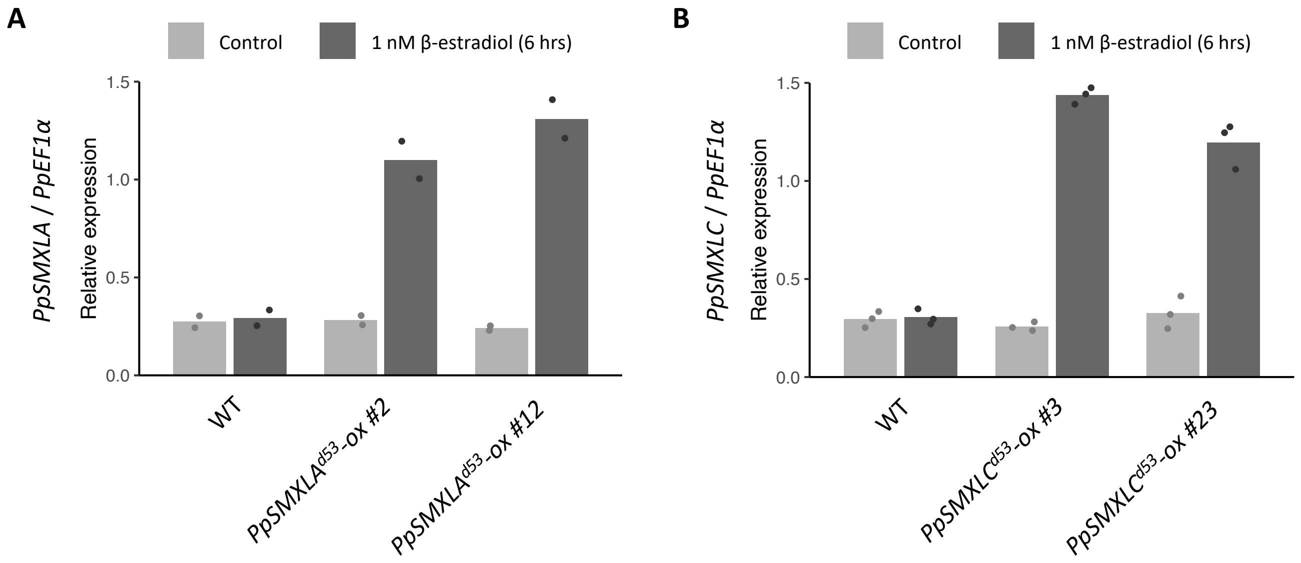


**Fig. S2. Induction of *PpSMXLA$^{d53}$* and *PpSMXLC$^{d53}$* over-expression.**

(A, B) Expression level of *PpSMXLA* in *PpSMXLA$^{d53}$-ox* lines (A) and *PpSMXLC* in *PpSMXLC$^{d53}$-ox* lines (B) after 1 nM β-estradiol treatment, examined by qRT-PCR (from 2 biological replicates in *PpSMXLA* and 3 biological replicates in *PpSMXLC*). *PpEF1α* was used as a reference gene.

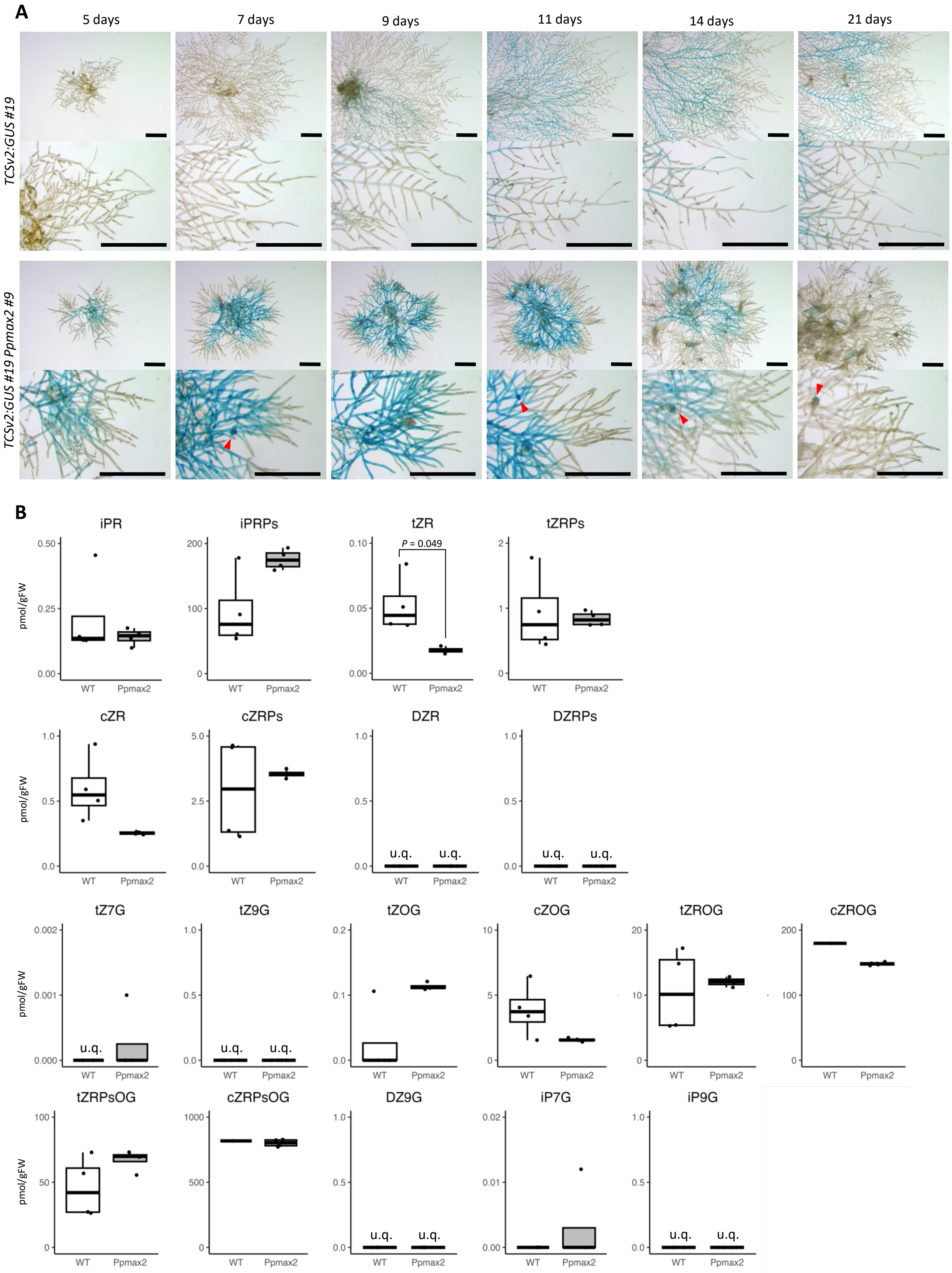


**Fig. S3. Analyses of cytokinins in WT and *Ppmax2* mutants.**

(A) Expression patterns of *TSCv2:GUS* in WT and a *Ppmax2* mutant during plant colony growth. Lower panels in each genotype show tip region of protonemata. Red arrowheads indicate initiating buds. (B) Quantification of cytokinins in WT and a *Ppmax2* mutant. "u.q." indicates under quantification limits. Statistical significance was evaluated by Student's *t* test (n=4). The *P* value is shown when it is statistically significant ($P < 0.05$). Scale bars, 500 μm.

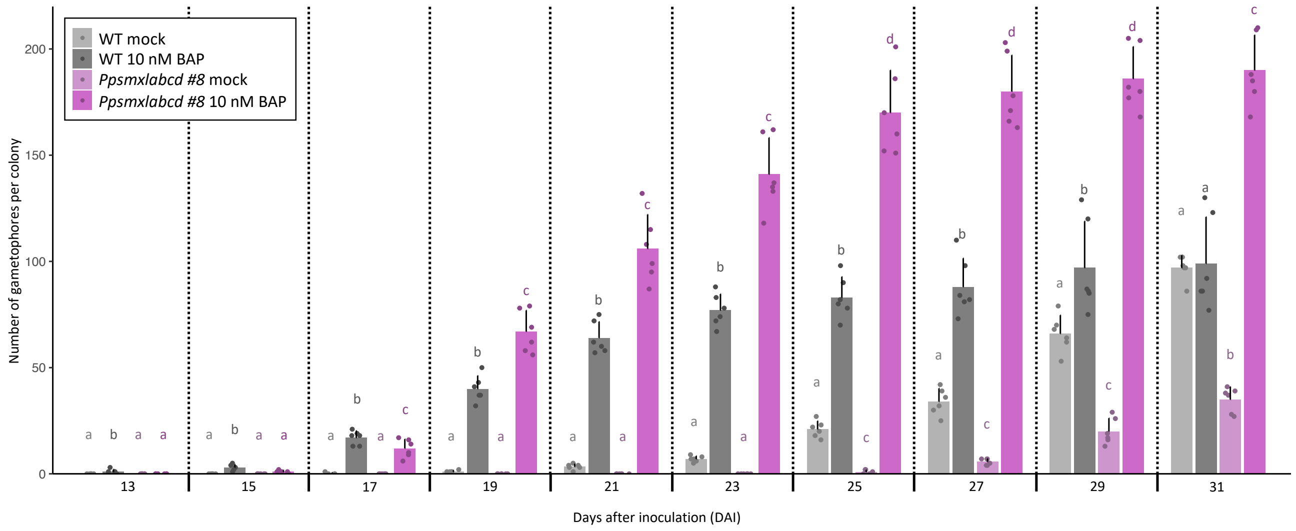


**Fig. S4. Effects of cytokinin in the gametophore formation.**

Number of gametophores per colony in WT and *Ppsmxlabcd* quadruple mutants grown on BCD medium supplemented with 10 nM BAP or solvent (Mock). Statistical significance was evaluated by HSD test (n=6, $p < 0.05$, statistical differences are indicated by different letters).

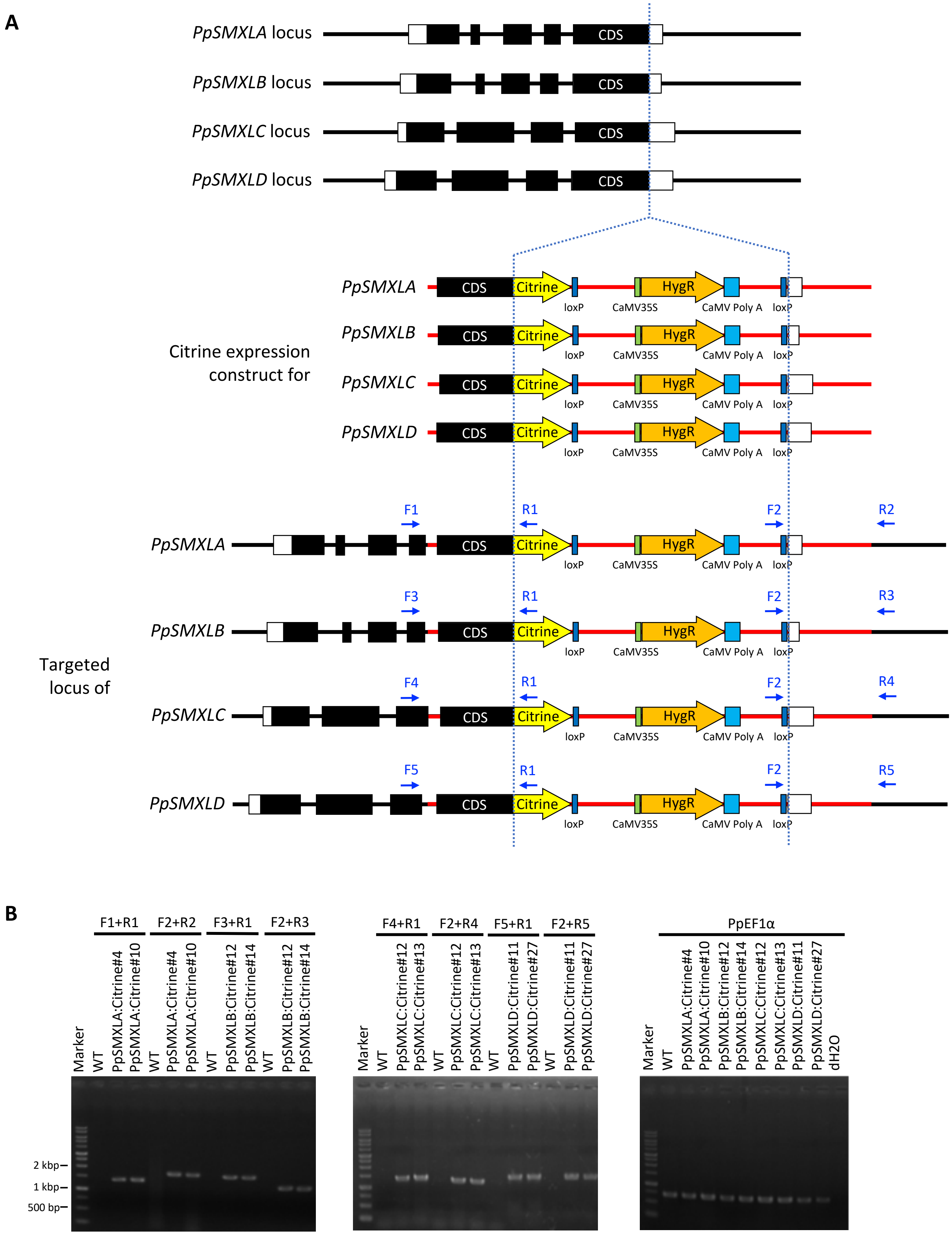


**Fig. S5. Construction for *PpSMXL:Citrine* lines.**

(A) Schematic diagrams showing DNA fragment introduced to *PpSMXL*s locus to visualize protein localizations of PpSMXLs. (B) Genotyping results to check the integration of the construct to the *PpSMXL*s locus through homologous recombination.

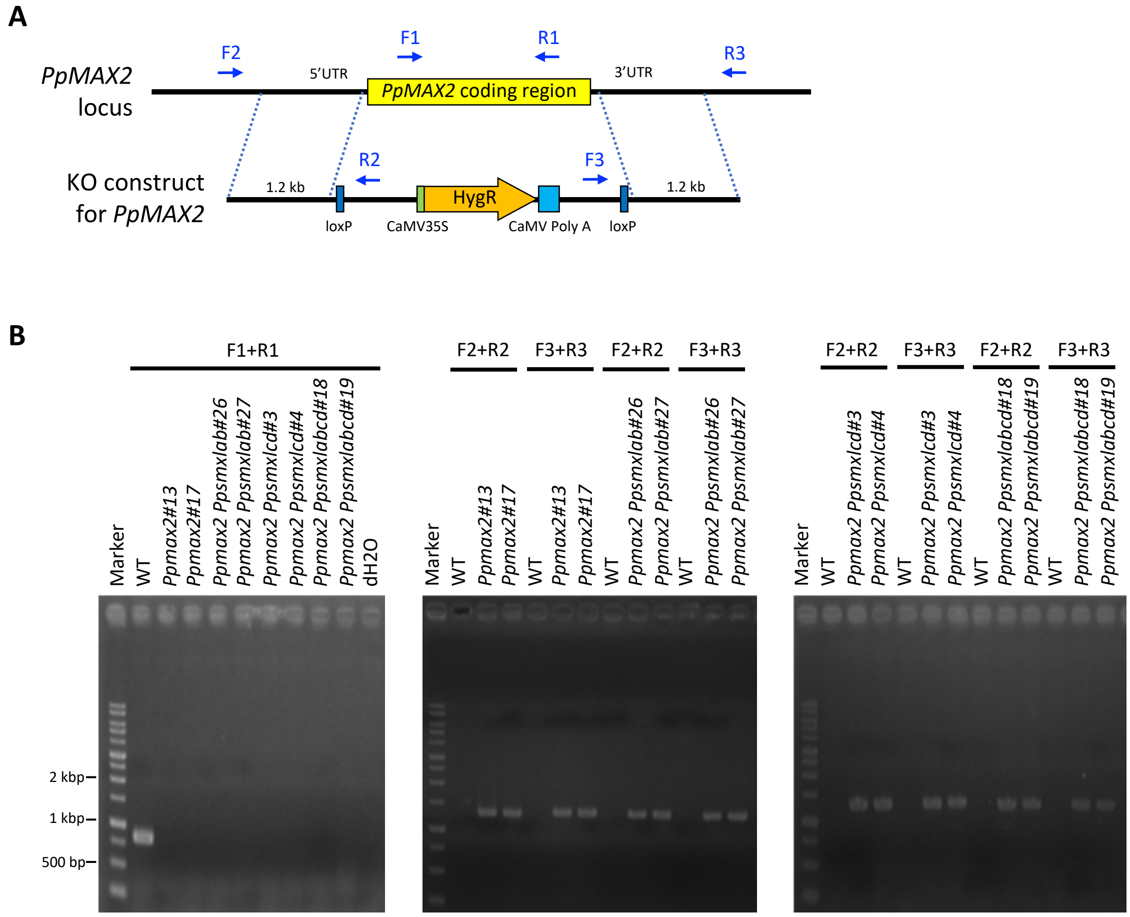


**Fig. S6. Construction for *Ppmax2* mutants.**

(A) Schematic diagrams showing DNA fragment introduced to the *PpMAX2* gene locus to produce *Ppmax2* mutants. (B) Genotyping results to check the loss of *PpMAX2* coding sequence and integration of the construct to the *PpMAX2* locus through homologous recombination.

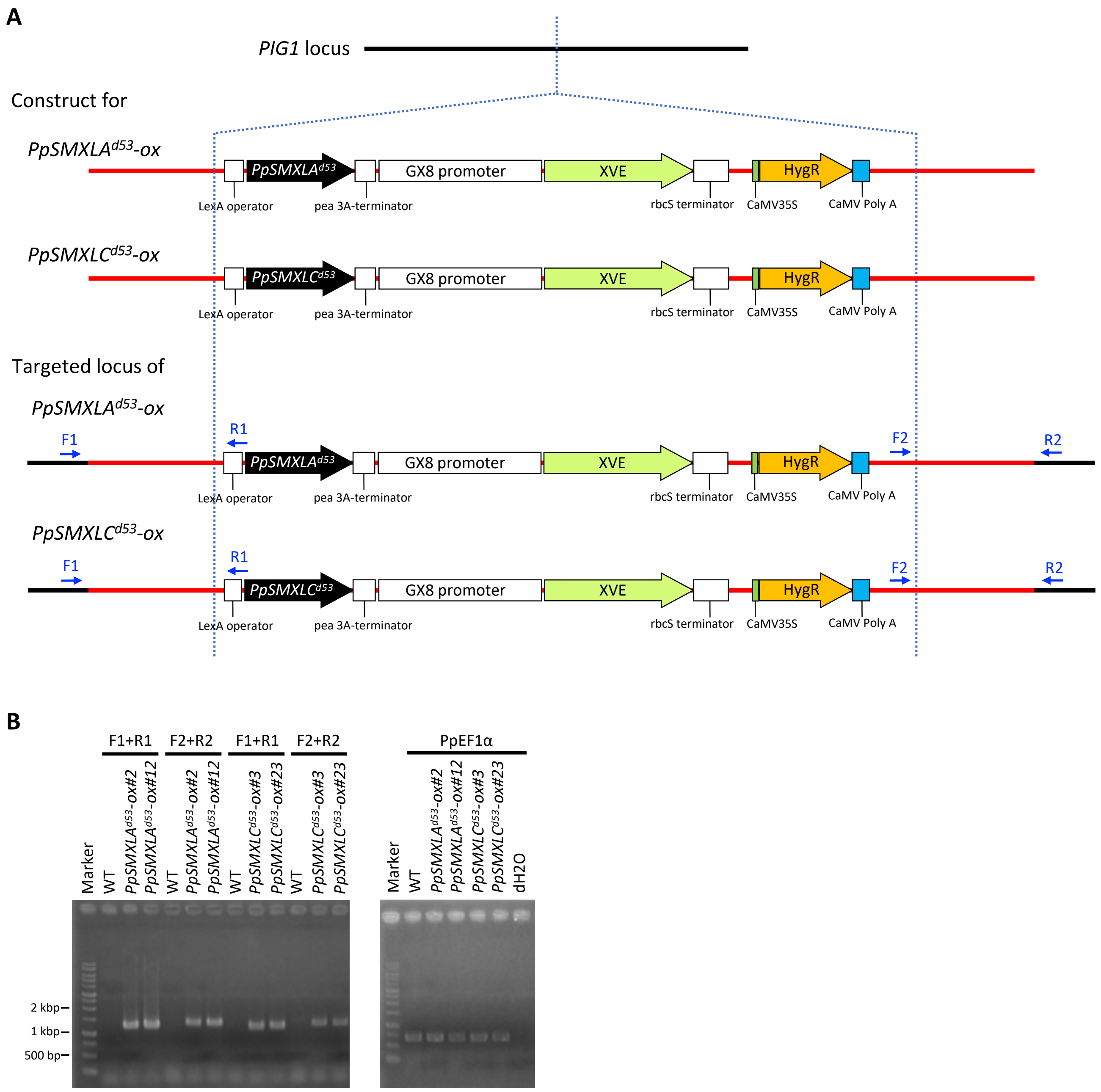


**Fig. S7. Construction for *PpSMXLA$^{d53}$-ox* and *PpSMXLC$^{d53}$-ox* lines.**

(A) Schematic diagrams showing DNA fragment to generate *PpSMXLA$^{d53}$-ox* and *PpSMXLC$^{d53}$-ox* lines. (B) Genotyping results to check the integration of the construct to the *PIG1* locus through homologous recombination.

**A**

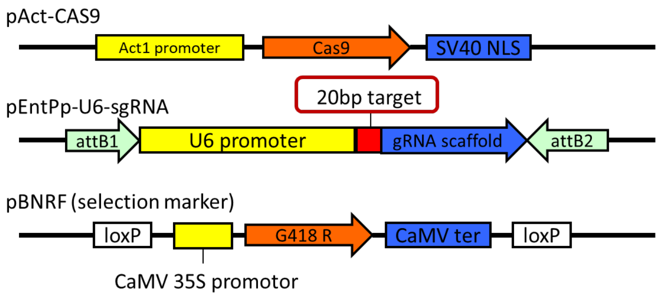


**B**

| | target sequence | PAM | |
|---|---|---|---|
| *PpSMXLA* : | GAATCAACCGCTGGTGATCT | TGG | |
| gRNA : | GAATCAACCGCTGGTGATCT | | (20bp) |
| *PpSMXLB* : | GAATCAACAGTTTGCGACCT | TGG | |
| gRNA : | GAATCAACAGTTTGCGACCT | | (20bp) |
| *PpSMXLC* : | GTTACCGAGGCTCGGAGGAG | GGG | |
| gRNA : | GTTACCGAGGCTCGGAGGAG | | (20bp) |
| *PpSMXLD* : | GTCACCGAGGCACGGAGAAG | GGG | |
| gRNA : | GTCACCGAGGCACGGAGAAG | | (20bp) |

**C**

*PpSMXLA*

```
WT 748          GAATCAACCGCTGGTGA--------------------------TCTTGGAACGACCTCTG  791
Ppsmxlab_#1     GAATCAACCGCTGGTGA-------------------------TTCTTGGAACGACCTCTG
Ppsmxlab_#2     GAATCAACCGCTGGTGAAACCTTCTTTCGTTCTAAAGGTTTATTCTTGGAACGACCTCTG
Ppsmxlab_#10    GAATCAACCGCTG----------------------------------------------G
Ppsmxlabcd_#8   GAAT---------------------------------------TCTTGGAACGACCTCTG
Ppsmxlabcd_#13  GAATCAACCGCTGGT----------------------------TCTTGGAACGACCTCTG
```

*PpSMXLB*

```
WT 998          GGAATCAACAGTTTGCGA-CCTTGGAAAAACCTTCTTTCG  1036
Ppsmxlab_#1     GGAATCAACAGTTTGCGAACCTTGGAAAAACCTTCTTTCG
Ppsmxlab_#2     GGAATCAACAGTTTGCGA--------AAAACCTTCTTTCG
Ppsmxlab_#10    GGAATCAACAGTTT-----CCTTGGAAAAACCTTCTTTCG
Ppsmxlabcd_#8   GGAATCAACAGTTTGCGATCCTTGGAAAAACCTTCTTTCG
Ppsmxlabcd_#13  GGAATCAACAGTTTGCGATCCTTGGAAAAACCTTCTTTCG
```

*PpSMXLC*

```
WT 61           AAGTATGCGGTTACCGAGGCTCGGAG-------GAGGGGTCACCCTCAGGTGCAACCCCT  113
Ppsmxlcd_#6     AAGTATGCGGTTACC-----------------------------CTCAGGTGCAACCCCT
Ppsmxlcd_#7     AAGTATGCGGTTACCTGAG------------------GGTCACCCTCAGGTGCAACCCCT
Ppsmxlcd_#12    AAGTATGCGGTTACCGAGGCTCGGAGACAACATGGAGGGGTCCCCTCAGGTGCAACCCCT
Ppsmxlabcd_#8   AAGTATGCGGTTACCGAGGCTCGGTA-----------------CCTCAGGTGCAACCCCT
Ppsmxlabcd_#13  AAGTATGCGGTTACCGAGGCTCGGTA-----------------CCTCAGGTGCAACCCCT
```

*PpSMXLD*

```
WT 61           AAGCATGCGGTCACCGAGGCACGGAGAAGGGGCCACCCCCAGGTGCAACC  110
Ppsmxlcd_#6     AAGCATGCGGTCACCGAGGCAC--------------CCCCAGGTGCAACC
Ppsmxlcd_#7     AAGCATGCGGTCACCGAG--------------------------GCAACC
Ppsmxlcd_#12    AAGCATGCGGCCACC----------------------CCCAGGTGCAACC
Ppsmxlabcd_#8   AAGCATGCGGTCACCGAGGCAC--------------CCCCAGGTGCAACC
Ppsmxlabcd_#13  AAGCATGCGGTCACCGAGGCAC--------------CCCCAGGTGCAACC
```

**Fig. S8. Construction for *Ppsmxls* mutants.**

(A) Schematic diagrams of constructs to generate *Ppsmxl*s mutants by CRISPR/Cas9 system. (B) DNA sequences of target sites for gRNA. (C) Mutation pattern of *Ppsmxl*s mutants generated by the CRISPR/Cas9 system examined by the Sanger sequencing. All mutations cause frame-shift leading to premature arrest by a stop codon. Numbers indicated in the WT sequences refer to the position in the coding sequence relative to the start codon.

**Table S1. List of primers used in this study.**

| Primer name | Sequence (5´→3´) | Project |
|---|---|---|
| S_A_end_F | ATCAAGTCGGAAAGAGGCTT | |
| S_A_end_R | TGATCCTCCTCCACTGCAGGCAACTTCAAT | |
| S_A_3UTR_F | AGTTTTCCTTTTCTTCTTCTTTTC | |
| S_A_3UTR_R | GATATCCATGAATGGATGGAAC | |
| S_B_end_F | ATCAATAGTGAATGCACAAACTG | |
| S_B_end_R | TGATCCTCCTCCACTGCAGCCTATTTCGAT | |
| S_B_3UTR_F | AGTTCTATTTATTTGCCTTTCT | |
| S_B_3UTR_R | GATATCCGGATAGTGGAAACC | Homologous region cloning of *PpSMXL:Citrine* reporter construct by homologous recombination |
| S_C_end_F | ATCGCCTGCAAGGTTGGAA | |
| S_C_end_R | TGATCCTCCTCCCACAGCCACGGTGGAC | |
| S_C_3UTR_F | CATCAAGGTCTTTATAGAACAG | |
| S_C_3UTR_R | GATATCTATTATTGAGAAGGTGCT | |
| S_D_end_F | ATCCCACTTTGAAAGGCCTT | |
| S_D_end_R | TGATCCTCCTCCCACAGCCATGGTGGAC | |
| S_D_3UTR_F | GTTAAAGGTCTTTACAGGC | |
| S_D_3UTR_R | AGGCCTATTGTGACAAGAATT | |
| Crispr_sgRNA_PpSMXLA_F | CTGGTGATCTGTTTTAGAGCTAGAAATAGCAAGTT | |
| Crispr_sgRNA_PpSMXLA_R | CGGTTGATTCATGGGTTGTAAGTCCTCCACCT | |
| Crispr_sgRNA_PpSMXLB_F | TTTGCGACCTGTTTTAGAGCTAGAAATAGCAAGTT | |
| Crispr_sgRNA_PpSMXLB_R | CTGTTGATTCATGGGTTGTAAGTCCTCCACCT | CRISPR guide RNA vector pENPp-U6-sgRNA site-directed mutagenesis inverse PCR |
| Crispr_sgRNA_PpSMXLC_F | CTCGGAGGAGGTTTTAGAGCTAGAAATAGCAAGTT | |
| Crispr_sgRNA_PpSMXLC_R | CCTCGGTAACATGGGTTGTAAGTCCTCCACCT | |
| Crispr_sgRNA_PpSMXLD_F | CACGGAGAAGGTTTTAGAGCTAGAAATAGCAAGTT | |
| Crispr_sgRNA_PpSMXLD_R | CCTCGGTGACATGGGTTGTAAGTCCTCCACCT | |
| PpMAX2_5'UTR_CLO_F | GGTGGTTTGCTCCGTTTTCC | |
| PpMAX2_5'UTR_CLO_R | ACAATGAGGACGCAGCAAGA | Homologous region cloning of *PpMAX2* gene disruption construct by homologous recombination |
| PpMAX2_3'UTR_CLO_F | TCTATGAGCTGGTGCGTGTG | |
| PpMAX2_3'UTR_CLO_R | GGCGAAAATTGTGGTGGCAT | |
| SMXLA_cDNA_pcr_F | CACCATGCGCTCTGGAGCTG | |
| SMXLA_cDNA_pcr_R | TTAACTGCAGGCAACTTCAATTTGA | Topo cloning of the coding regions of *PpSMXLA* and *PpSMXLC* |
| SMXLC_cDNA_pcr_F | CACCATGCGTAGCGGGGCAAATTCAGTCC | |
| SMXLC_cDNA_pcr_R | CTACACAGCCACGGTGGACA | |
| SMXLA_endcheck_F | TGGAGAAGGTTGTGTGGCAG | |
| SMXLA_3utrcheck_R | TGGTGAATGTAGATGAGTGGGC | |
| SMXLB_3utrcheck_R | GGAATTGCAACAACACAGGCT | |
| SMXLB_endcheck_F | CACAAACCCGAGCCCCTTAT | |
| SMXLC_endcheck_F | TCATGAAGTGTCGGTCTGGC | |
| SMXLC_3utrcheck_R | ACAAGATCCACAATAAAGAACTTGTG | Genotyping of *PpSMXL:Citrine* reporter lines |
| SMXLD_endcheck_F | GGGGGCTTATGTGGACTCTC | |
| SMXLD_3utrcheck_R | ACTTTGATTATAGACAACAATGAAGGA | |
| HRCitSmaIseq.ck_F | ACCTCACCAAAATACGAAAGAGT | |
| HRCitEcoRVseq.ck_R | CTCGCCGGACACGCTGAA | |
| Crispr_PpSMXLA_check_F | AGGCATGTGCTGATACGCAT | |
| Crispr_PpSMXLA_check_R | ACTGCTTTGTTTTCGAACGGT | |
| Crispr_PpSMXLB_check_F | AGTGTCTCACCTTTTGGCGT | |
| Crispr_PpSMXLB_check_R | TAGCAGACTGGGTCCTCCAT | Genotyping of *Ppsmxls* |
| Crispr_PpSMXLC_check_F | CGGGGCAAATTCAGTCCAAC | |
| Crispr_PpSMXLC_check_R | TGACATTCTTGATCCTGGGCC | |
| Crispr_PpSMXLD_check_F | CGTAGCGGGGCAAATTCAGT | |
| Crispr_PpSMXLD_check_R | TGTTCAAAACATCCTCGTCCCT | |
| PpMAX2_EcoRV_check_F | CGTGTTTTCCTTGGTTGCGT | |
| pTN186_EcoRV_seq.check.R | GGGGGCTGCAGGAATATAACT | Genotyping of *Ppmax2* |
| pTN186_SmaI_seq.check.F | AGTTATCCCTCACACCGGTG | |
| PpMAX2_SmaI_check_R | TCATCACGTGCCACCTTTGT | |
| HR_pPGX8_FAU5'geno_F | GACTTGTGCCCTGAATGTAC | |
| HR_pPGX8_FAU5'geno_R | GTACGTCGAGGGGATGATAAT | Genotyping of *$PpSMXLA^{d53}$-ox* and *$PpSMXLC^{d53}$-ox* |
| HR_pPGX8_FAU3'geno_F | GTGCAAGGTAAGAAGATGGAAA | |
| HR_pPGX8_FAU3'geno_R | AATCTGGGAATAGCTTGTTATTGT | |
| qPCRforPpEF1a_F | ACTAGGGAGCACGCGTTGTT | |
| qPCRforPpEF1a_R | TACTTCGGGGTGGTTGCGTC | |
| SMXLA_q-pcr_F2 | GTGAACCACCTTAGTCAGAGGC | |
| SMXLA_q-pcr_R2 | ACTCTTGAGTAGGTGCGTTTCC | RT-qPCR analysis |
| SMXLC_q-pcr_F | CGGGAAAGGGTTGACTAGCC | |
| SMXLC_q-pcr_R | TCATTCTCCTCTTGCGTCACG | |
| qPCRforPpCKX1_F2 | GGCGGCACTCTATCGAATGC | |
| qPCRforPpCKX1_R2 | GTAGGCGTGCATTGTACCACC | |
